# Full Likelihood Inference from the Site Frequency Spectrum based on the Optimal Tree Resolution

**DOI:** 10.1101/181412

**Authors:** Raazesh Sainudiin, Amandine Véber

## Abstract

We develop a novel importance sampler to compute the full likelihood function of a demographic or structural scenario given the site frequency spectrum (SFS) at a locus free of intra-locus recombination. This sampler, instead of representing the hidden genealogy of a sample of individuals by a labelled binary tree, uses the minimal level of information about such a tree that is needed for the likelihood of the SFS and thus takes advantage of the huge reduction in the size of the state space that needs to be integrated. We assume that the population may have demographically changed and may be non-panmictically structured, as reflected by the branch lengths and the topology of the genealogical tree of the sample, respectively. We also assume that mutations conform to the infinitely-many-sites model. We achieve this by a controlled Markov process that generates ‘particles’ in the hidden space of SFS histories which are always compatible with the observed SFS.

To produce the particles, we use Aldous’ Beta-splitting model for a one parameter family of prior distributions over genealogical topologies or shapes (including that of the Kingman coalescent) and allow the branch lengths or epoch times to have a parametric family of priors specified by a model of demography (including exponential growth and bottleneck models). Assuming independence across unlinked loci, we can estimate the likelihood of a population scenario based on a large collection of independent SFS by an importance sampling scheme, using the (unconditional) distribution of the genealogies under this scenario when the latter is available. When it is not available, we instead compute the joint likelihood of the tree balance parameter *β* assuming that the tree topology follows Aldous’ Beta-splitting model, and of the demographic scenario determining the distribution of the inter-coalescence times or epoch times in the genealogy of a sample, in order to at least distinguish different equivalence classes of population scenarios leading to different tree balances and epoch times. Simulation studies are conducted to demonstrate the capabilities of the approach with publicly available code.

## 1 Introduction

Demographic inference based on genetic data has been a major challenge in the last two decades. Many methods and algorithms have been developed to turn the genetic diversity observed in a sample of individuals into reliable estimates of population structure or demography. Most of them consist of decomposing the question into (*i*) computing a weight or likelihood function for the parameters of interest under a given genealogical tree of the sample, and (*ii*) aggregating these weights by averaging over the set of all possible genealogies to account for the fact that they are hidden in practice. However, even for moderately large sample sizes (*n* ≥ 5 or 10), the size of the state space of the full genealogy is huge and such an averaging cannot be performed in an exact way. Instead, one resorts to exploring the set of labelled trees as exhaustively as possible, e.g. via Monte Carlo methods, but the associated computational cost grows extremely quickly with the sample size. In this work, we exploit the fact that the distribution of the *Site Frequency Spectrum* (SFS) depends on a coarser description of the genealogical tree of the sample than the classical leaf-labelled binary tree representation, which significantly decreases the size of the space to explore when computing likelihoods.

Let us consider a locus which is free of intra-locus recombination, at which mutation occurs at rate *θ*. We sample n individuals at present and look at how mutations are shared among them. Under the infinitely-many-sites model, these Poissonian mutations give rise to a site frequency spectrum *S* = (*S*_1_,…, *S*_*n*-1_), which reports how many mutations are carried by each given number of individuals in the sample:

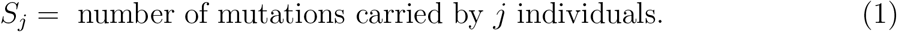

The site frequency spectrum is used routinely in population genetics inference. See [GJB13] and references therein for a recent review.

SFS carries considerable information about the underlying hidden genealogical tree with mutations upon it. In this paper, we introduce a novel importance sampler, which produces a *tree topology matrix F*, a *mutation pattern matrix M* on the tree, and a vector of *epoch times T* = (*T*_2_,…,*T_n_*), such that *F*, *M* and *T* are compatible with the given SFS *S*, i.e. 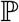(*S* | *F*, *M*, *T*) > 0. Here, the epoch time *T_k_* is the amount of time during which the sample has exactly *k* ancestors. When the distribution (not conditioned on the observed SFS) of the hidden genealogy of a sample under the family of population scenarios of interest is known, we can use these triplets to estimate the corresponding likelihood function *given the observed SFS*. In particular, setting *β* = 0 in our procedure enables us to compute the likelihood of a given scenario in a (parametric or not) family of stochastic past population size trajectories (*N_t_*, *t* ≥ 0), assuming that the genealogy of a sample is given by the Kingman coalescent with fluctuating population size. See Section 3 for some parametric examples, including exponential growth and bottlenecks. If the unconditional law of the sample genealogy is not known, we can instead resort to supposing that the tree topology is well-described by Aldous’ Beta splitting model (see below), which specifies a measure on tree balance through a single parameter *β*, and then estimate the joint likelihood of the parameter *β* and of a demographic model dictating the law of the epoch times. Under a given model of demography specifying the distribution of epoch times, our procedure would then allow one to distinguish between different equivalence classes of historical population scenarios up to their tree topology distributions specified by different values of *β*. Although we illustrate how to apply such a *Beta-splitting demographic model* in a semi-parametric spirit upon data simulated from a complicated scenario using `ms` [Hud02] in Section 3.5, we emphasize that it is not the point of this paper to study the effect of any given structural scenario on the β-specific shapes of the genealogical trees, or to inquire which scenarios can be distinguished based on their balance parameters.

The main strengths of our approach reside in the following points. First, we use the minimal tree resolution on which the law of the SFS depends, in the sense of [SSV15]. More precisely, instead of the set of all leaf-labelled binary tree topologies (on which the Kingman coalescent is defined), we work with the set of all binary tree shapes (also called the *unvintaged and sized* tree resolution) encoded by our tree topology matrix (*F_k,j_*)*_k,j_* which only tracks the number of edges in the *k*-th epoch of the tree that subtend j leaves through *F_k, j_*. See below for a more precise definition. This drastically reduces the state space of our sampler. Indeed, by Proposition 12 in [SSV15], each binary tree shape represents of the order of *n*!/2^*n*-1^ (at the very least) leaf-labelled binary tree topologies; as an illustration, note that *n*!/2^*n*-1^ is approximately 7100 for *n* = 10, and 4.6 × 10^12^ for *n* = 20. Furthermore, only a subset of these shapes is compatible with the SFS and our sampler is able to produce only compatible tree shapes, each such shape having a positive probability of being produced (that is, no compatible shape is potentially missed by the sampler). The significant reduction of the state space of tree topologies enables us to separately consider SFS-compatible genealogical histories at each one of a number of independent non-recombining loci — a natural framework for subsequent locus-specific outlier (non-neutrality) detection.

Second, the sampler needs to be informed of population-level processes equally affecting the genealogies at all independent loci only through (*i*) their tree topology or shape via the parameter *β* ∈ (–2, ∞) in Aldous’ Beta-splitting model [Ald01], introduced below to account for topologically sensitive processes including certain types of population structure and sampling schemes, and (*ii*) their branch lengths via a vector of *a priori* mean epoch times which typically summarize the demographic scenario experienced by the population. In fact, one major difference between the signature left by population structure and that left by fluctuating population sizes lies in the balance of the genealogical tree topologies, which has been barely explored even in the classical models. This characteristic balance of the genealogies influences the observed mutation pattern and could thus be at the basis of a straight-forward test for population structure affecting all loci or non-neutral evolution affecting only loci under selection or linked to a recently selected site. The importance sampler and likelihood procedure developed in this paper allow not only to test for deviations from panmixia and neutrality (corresponding to *β* = 0 in what follows), but also to infer the most likely balance parameter *β* corresponding to a given SFS, or set of SFSs from independent loci representing processes that affect the whole population.

As described above, the sampling of a ‘particle’ (*F*, *M*, *T*) uses the Beta-splitting model of tree balance in the topology matrix *F*, and only requires some *a priori* demographic laws for epoch times in *T*. The expected epoch times under these demographic laws can be obtained either analytically, or by an easy round of simulations from *any* standard demographic model, including parametric models, such as exponential growth or bottleneck, or semi-parametric models involving the class of piecewise constant or exponential functions, for example.

A rather large panel of methods already exist to infer demographic parameters from the observed SFS, but none of them are able to compute the full likelihood of a given SFS at a non-recombining locus. The *Poisson Random Field* approach [SH92, Nie00, GHWB09] considers a series of independent SNPs in a sample of size *n*. Assuming the infinitely-many-sites mutation model with a very low mutation rate, the distribution of the number of mutations (or derived alleles) carried by *k* ∈ {1,…, *n* – 1} individuals is either approximated by a Poisson distribution whose parameter is given by the ratio of the average length of all the edges in the genealogical tree that subtend *k* leaves to the average total length of the tree; or it is described as a Poisson random variable whose parameter is given by the probability that *k* out of *n* individuals are sampled within the current population carrying the derived allele. The average length of edges subtending a given number of sampled individuals is usually obtained by simulation as in [Nie00], while the sampling probability is obtained by solving a Wright-Fisher diffusion with selection as in [GHWB09]. Our procedure generalizes this approach. Indeed, the Poisson random field methodology imposes that each locus should be a single segregating site that is at an infinite recombinational distance from all other segregating sites, and thereby misses the shared genealogical signal at linked sites. In contrast, the likelihood that we compute does not assume that all SNPs are independent at the locus of interest, nor does it assume that the probability of a given mutation being carried by exactly *k* individuals is the *average* (over many tree realizations) of the proportion of the tree length corresponding to the total length of *k*-edges. Instead, our procedure can handle SFS data with fully-linked SNPs within each given locus, by constructing locus-specific particles whose genealogical and mutational histories are compatible with the SFS at the locus. We can then extend it to several independent loci which are free of intra-locus recombination to reduce the variance of the likelihood-based estimator. Now, assuming that the mutation rate is very small so that we see at most one mutation per locus, the recombinational distance between adjacent loci is very large, and the tree balance parameter *β* is 0 in order to enforce the Kingman coalescent prior on tree topologies, our sampler is essentially equivalent to the Poisson Random Field approach of [Nie00], with the notable difference that the probability of seeing a mutation carried by *k* individuals is now computed from the true probability of the placement of the mutation *conditionally on the tree topology*, for every particle generated by the sampler for each locus with possibly more than one segregating site.

In [BFL15], a method based on the probability generating function of the branch lengths in the genealogies is developed to extract the signal in SFS from linked sites and applied to detect the occurrence of a bottleneck in the history of the population, relying again on the Kingman coalescent model. Despite the use of unlabelled tree shapes and other clever tricks to take advantage of the symmetries of the problem, deriving the probability of a given SFS requires the sum over a very large set of mutation placements on the tree which are compatible with the SFS, a generally unfeasible computation whenever *n* > 5.

Skyline plots form another family of inference methods for demographic history, as reviewed in [HS11]. These nonparametric methods rest on the assumption that there is not much variability in the SFS-compatible reconstructed tree on which the estimation of the local harmonic means of effective population size is based (c.f., [PRH00]). However, this will typically not be the case when the individual mutation rate is low and the SFS contains only a few mutations. [HD08] extend the method to several unlinked loci, enhancing the demographic signal captured by the reconstructed trees at the different loci.

All these approaches assume independent loci free of intra-locus recombination, which may be sensible if we consider short loci far enough from each other in the genome. Following the improvement of the accessibility of whole-genome data and of the mathematical modelling of linkage, different methods focusing on large stretches of recombining DNA have been set up and used to reconstruct population histories. For instance, [HN13] study the set of distances between neighbouring SNPs within a long sequence of genome from a sample of two individuals. They derive an approximate formula for the distribution of the typical length of a tract of identity by state (IBS) using the Sequentially Markov Coalescent (SMC) of [MC05] and the related SMC’ model of [MW06]. Assuming these pieces of IBS sequences are nearly independent, they use a composite likelihood approach to infer the parameters in a model incorporating population size changes, and divergence and admixture events between sub-populations. The same approach is used by [BEKV13] to reconstruct the lineage diffusion coefficient and the neighbourhood size in a spatially structured neutral population. The SMC or SMC’ approximation is also a key ingredient in the conditional haplotype sampling distribution developed by [SKS16] for population scenarios comprising discrete sub-populations related through migration and with potentially varying effective population sizes (described by a given class of functions, such as piecewise linear or piecewise exponential). [PWR15] model the effective population size of a population as the exponential of a Gaussian process. Assuming that the local genealogies follow the SMC’ model, the pattern of diversity observed in the sequence data is used to reconstruct the fluctuations of the effective population size in a Bayesian nonparametric way. Note that the choice made there to encode the tree topologies at the sufficient resolution of the *ranked tree shapes* [Taj83, SSV15] considerably enhances the exploration of this component of the state space during the MCMC step, a point which is pushed further by [PVWR18]. Such methods explicitly try to model recombination within a locus, involving a very large hidden space of compatible histories when compared to recombination-free histories. Our approach generalizes the Poisson random field methods and can be complementary to methods that model recombination explicitly when applied to recombinationally distant blocks of contiguous sites which are devoid of any signal of recombination within the block. This assumption, along with our parametric model for tree balance, allows us to extract information from SFS in a locus-specific manner across thousands of loci that are recombinationally independent for demographic inference and outlier detection.

To overcome the difficulty of computing analytical (or even approximate) likelihoods in potentially complex population models, a simple simulation-intensive approach known as Approximate Bayesian Computation or ABC [BZB02] is now routinely used in a wealth of studies. Recently [BRJ+16] demonstrate through simulations that considering the joint information provided by SFS and linkage disequilibrium improves the accuracy of parameter reconstruction when compared to a method based only on SFS or LD. An ABC methodology is used by [PWE10] to distinguish different demographic and structural scenarios using a variety of statistics of microsatellite data. A significant disadvantage of any ABC method is the lack of locus-specific likelihood for the SFS itself, the basic quantity that the method tries to circumnavigate from computing. Typically, the SFS data across multiple loci is reduced to the mean and variance of various *ad hoc* summary statistics of the SFS across all the loci during the simulation-intensive approximations underlying ABC and thus no efforts are made to integrate over hidden genealogical histories that are compatible with the SFS at each locus. In contrast, our approach develops a controlled Markov process to obtain the likelihood of each SFS directly. As described below, this controlled Markov process is the step-by-step construction of a tree topology and a vector of epoch times, controlled by a vector of presence or absence of mutations carried by each number of individuals. This vector is used to ensure that at every step *k* ≥ 1, (at least) one edge subtending *n* – *k* leaves is necessarily placed in the tree whenever the component *S_n-k_* of the SFS is positive. The resulting tree is thus always compatible with the given SFS.

The methods reviewed here are well-suited for inferring demographic parameters that affect all loci, since they use the combined information coming from the per-locus site frequency spectra generated by their common history. In a few instances, they have also been used to detect outlier loci potentially subject to natural selection [Nie05, RPR_+_12]. However, when one wants to infer the locus-specific history provided by the SFS, to our knowledge there are no known methods analogous to the importance samplers available for inference from the full *binary incidence matrix* or BIM data [FD01, DIG04, HUW08, KJS15, KSCS16], except the naive importance sampler via controlled Markov chains in [STH+11]. Here we propose an efficient importance sampler that uses very natural *a priori* information on the law of hidden genealogical histories to produce a triplet (*F*, *M*, *T*) at the minimally sufficient resolution of tree topology, mutation history and epoch times that are compatible with a given SFS. Since the hidden state space has been optimized, our method can cope with large numbers of samples and independent loci to obtain maximum likelihood estimates or MLEs of parameters in (*i*) demographic models with analytical expressions for likelihood, such as, exponential growth or bottleneck models (under Kingman coalescent with *β* = 0) and in (**ii**) Beta-splitting demographic models with *β* G (–2, ∞) under a simple demographic scenario specified by exponential rates for epoch times.

Such Beta-splitting demographic models are meant to be a simple semi-parametric approximation of an equivalence class of much more complex historical processes with no analytically tractable likelihood function, such as time-varying migration structures, admixture times and/or sub-population specific demographies. Such an approximation may be an acceptable alternative to simulation-intensive inference methods such as ABC that require one to envision specific high-dimensional parametric families of historical scenarios to simulate from. Furthermore, since the particle systems in the hidden space are constructed locus-specifically, the likelihood and MLEs at each locus can be used directly for outlier detection.

The paper is organized as follows. In Section 2, we progressively present our sampler by introducing the *a priori* laws used for the tree topology (§2.1) and the vector of epoch times (§2.2), then by precisely defining the triplet (*F*, *M*, *T*) describing a genealogical tree with mutations at the optimal resolution (§2.3), and finally by giving a full description of how the sampler produces such a ‘particle’ (§2.4). In Section 3, we perform some simulation studies for a few examples of Beta-splitting demographic models (Kingman’s coalescent with exponential growth in §3.2 and with bottlenecks in §3.4), and we estimate *β* for a complex population history simulated using `ms` in §3.5. Some generalizations of this approach to the inference of demographic and structural scenarios are discussed in Section 4. The proof-of-concept code for the sampler and the likelihood procedure as well as its integration with `ms` is publicly available at [SV18]. The detailed pseudo-code can be found in Section 1 of the Supplementary Material.

## 2 The Sampler

In all that follows, we assume that the genealogical tree relating a sample of **n** individuals is always binary. A coalescence event therefore corresponds to the number of ancestral lineages decreasing from some *k* ∈ {2,…, n} to *k* – 1. We call epoch *k* the interval of time in the past during which the sample has **k** distinct ancestors, and write *T_k_* for the duration of this epoch. In other words,

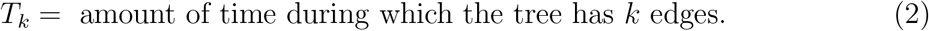

Mutations are assumed to occur at some per-lineage rate *θ* > 0 along the branches of the tree, and to conform to the infinitely-many-sites model. No recombination happens within the locus considered. See Figure 1 for an example of mutations on a stretch of nine sites which are completely linked.

**Figure 1:**
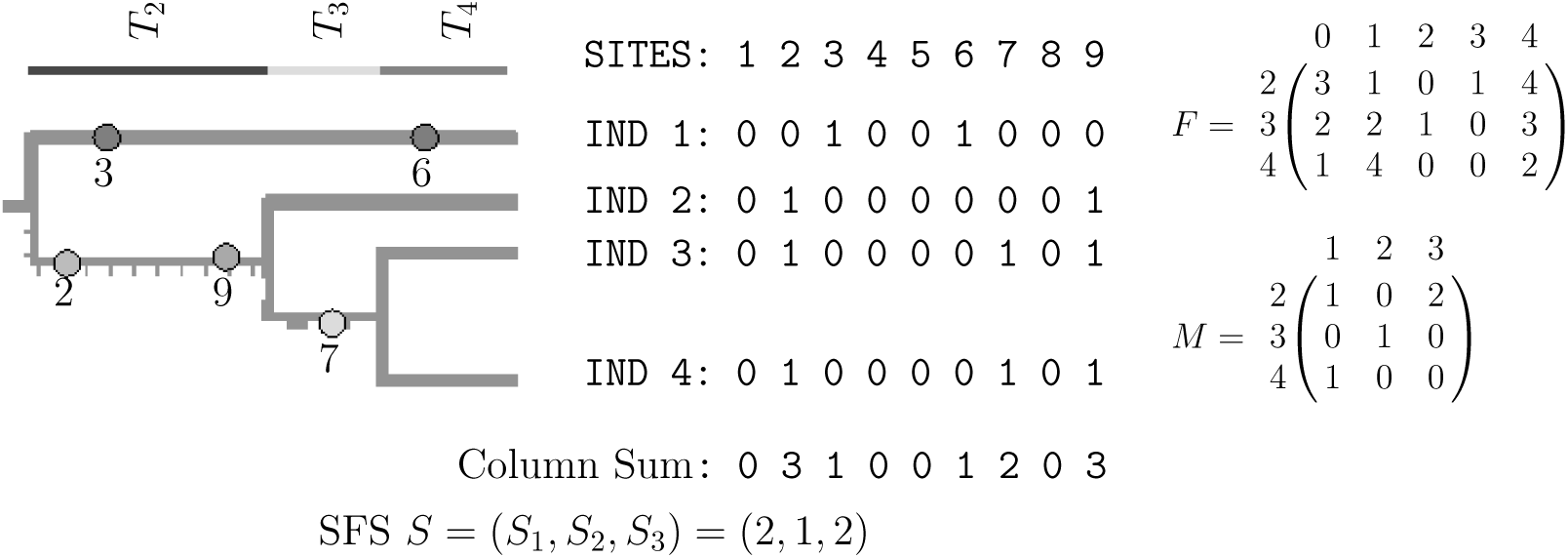
The coalescent tree with mutations on the three type of edges subtending 3, 2 and 1 leaves (left), the observed derived mutation incidence matrix with its site frequency spectrum *S* (middle) and the corresponding SFS history with topology matrix *F* and mutation matrix *M* (right). At most one mutation per site under the infinitely-many-sites model are superimposed as a homogeneous Poisson process upon the realization of identical coalescent trees at nine homologous sites labeled {1, 2,…, 9} that constitute a non-recombining locus from four individuals labeled {1, 2, 3, 4}.

Instead of integrating over the full history of leaf-labelled coalescent trees with mutations (as shown on the left side of Figure 1), our main idea here is to work with a much smaller space of topology matrices encoding the number of edges in epoch *k* that subtend *j* leaves, and mutation matrices recording the number of mutations that fall on such edges (as shown on the right side of Figure 1 by *F* and *M*, respectively). Explicitly, for every *k* ∈ {2,…, *n*} and *j* ∈ {1,…, *n* – 1},

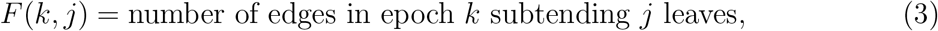

while

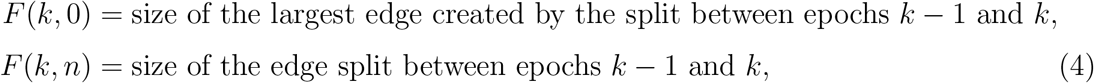

where the size of an edge corresponds to the number of leaves (or sampled individuals) that it subtends. The rows and columns of *F* thus range in 2, 3,…, *n* and 0,1,…, *n*, respectively, and necessarily *F*(*k*,*j*) = 0 if *j* ∈ {*n* – *k* + 2,…, *n* – 1}. Observe that the topology matrix *F* encodes more information than the *f*-sequence of [STH+11], the minimal sufficient topological statistic of the labelled genealogical tree of the sample (in addition to epoch times), that is necessary for the prescription of multinomial-Poisson probabilities for SFS (see Eqs. (6) and (7)). The extra information in *F* given by *F*(*k*, 0) and *F*(*k*, *n*) can in fact be derived from the other matrix entries, but are recorded in the two extra columns to ease the computations based on the *F*-matrix. Indeed, these coefficients are exactly what is needed to prescribe the topology probabilities dictating the tree shape under Aldous’ Beta-splitting model [Ald01], our simple parametric model of various phenomena affecting tree shape.

Similarly,

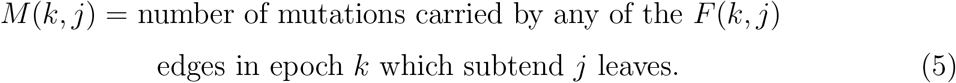

Therefore, the rows and columns of *M* range in 2, 3,…, *n* and 1,…, *n* – 1, respectively. We use standard notation for the sub-matrix of a matrix: *F*(*a* : *b*, *c* : *d*) is the sub-matrix of *F* made of rows *a*, *a* + 1,…, *b* and columns *c*, *c* + 1,…, *d*.

The product of the epoch time row vector *T* and the matrix *F*(2 : *n*, 1 : *n* – 1) gives the total length of the edges in the tree that subtend 1, *2,…, *n* –* 1 leaves as follows:

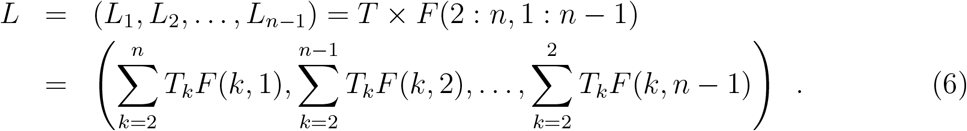

As shown in Proposition 5 of [STH+11], the probability of the SFS only depends on the coalescent tree through *L:* Conditionally on *L*,

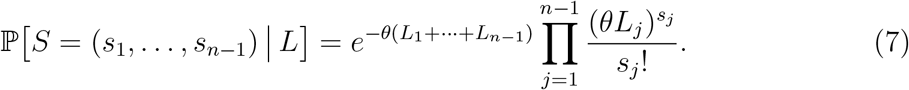

Thus, we only need to consider topology matrices and epoch time vectors to compute the likelihood of the observed SFS. However, in the incremental construction of *F* and T by our sampler we use *M* to take the partially constructed SFS history into account while ensuring that the fully reconstructed SFS history remains consistent with the observed SFS.

Next, we describe the sampler’s *a priori* laws for the tree topology and the epoch times.

### 2.1 *A Priori* Laws for the Topology

Our sampler uses a modification of the Beta-splitting model of [Ald01] which specifies the order of the splits. Aldous’ Beta-splitting model is a one-parameter family of random cladograms which has the advantage of containing several of the most classical null models for tree shapes used in phylogeny reconstruction ([MH97]), such as the *equal-rates-Markov* model (i.e., the random topology of the Kingman coalescent), the *proportional-to-distinguishable-arrangements* model or the *equiprobable-types* model. More precisely, for a given choice of *β* ∈ (–2, ∞), if an edge subtending *b* leaves (i.e., ancestral to *b* individuals in the sample) is split into two edges, then the probability that the two daughter edges subtend *x* and *b* – *x* leaves is given by

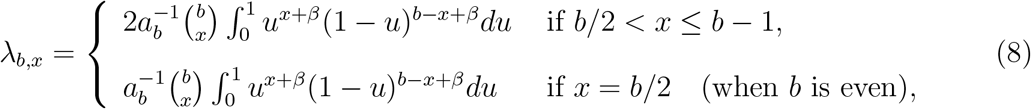

where *a_b_* is the normalizing factor

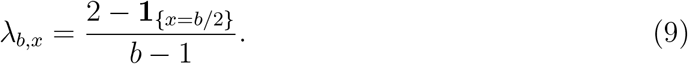

The particular case *β* = 0 corresponds to the topology of the Kingman coalescent, with

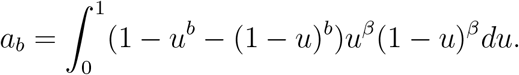

Choosing *β* close to –2 gives rise to comb-like trees, while *β* 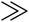 1 produces highly balanced tree topologies [Ald01]. Thus the Beta-splitting model gives us a one-parameter family spanning the whole range of possible tree balances.

Using (8), we can define the probability of producing a given topology *F* under our incremental Beta-splitting model. To simplify the notation, for every epoch *k* ∈ {2,…, *n*}, we write *m_k_* for the size (or number of leaves it subtends) of the edge split between epochs *k* – 1 and *k*, and *ℓ_k_* for the size of the largest edge created during this split. Recalling (4), we have *m_k_* = *F*(*k*, *n*) and *ℓ_k_* = *F*(*k*, 0).

We proceed by going from the root towards the leaves of the tree. First, the edge chosen to break at the beginning of epoch *k* ≥ 2 has size m with probability *F*(*k* – 1, *m*) * (*m* – 1)/(*n* – *k* + 1) (observe that *F*(1, ⋅) = (0,…,0,1), since epoch 1 formally corresponds to the period during which there is only one ancestor to the whole *n*-sample). The law of this random choice is arbitrary since it is not specified in Aldous’ *stick-breaking* construction ([Ald01]); it corresponds to the law of the choice of the block to be split at the beginning of the *k*-th epoch in the *forwards-in-time* unlabelled Kingman coalescent. Second, the split within the chosen block is given by the probability λ_*m*_ described in (8). Thus, we obtain that under the law 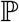_*β*_ of the incremental Beta-splitting model with parameter *β* ∈ (–2, ∞), the probability of a given topology *F* is given by

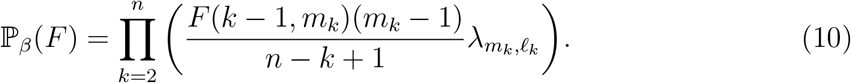

In particular, when *β* = 0, using (9) we see that if 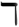(*F*) denotes the number of splits such that *ℓ_k_* ≠ *m_k_*/2, we indeed recover the probability

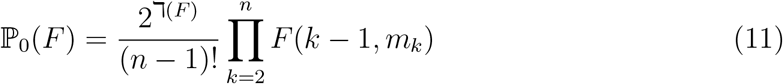

of the unvintaged and sized Kingman coalescent (see Proposition 11 in [SSV15]).

### 2.2 *A Priori* Laws for the Epoch Times

The epoch time component of our sampler is initialized with a sample from a vector *T*^0^ = (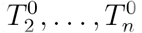) of *n* – 1 independent exponential random variables with respective parameters *A_k_*. The mean 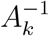 of 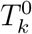 is taken to be the average length of the *k*-th epoch in the scenario whose likelihood we want to compute, independently of the observed SFS. For example, if we assume that the genealogy underlying the observed SFS conforms to the Kingman coalescent with effective population size *N*_0_, then

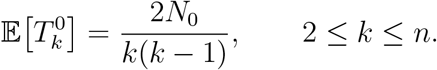

Other examples of parametric models are given in Section 3 of the Supplementary Material. The *a priori* rate vector (*A_k_*)_2≤k≤n_ is thus another input of the sampler. As mentioned in the introduction, its components can either be computed analytically in simple scenarios, or can be estimated by a round of simulations (without conditioning on the observed SFS).

The choice of exponential *a priori* laws for the epoch times is motivated by the following standard property of conditioned Gamma random variables: If *T* follows a Gamma distribution with parameters *k*, *λ*, denoted by *G*(*k*, *λ*) and with density

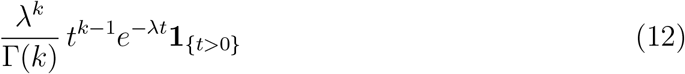

(where Γ is the Gamma function), then the law of *T* conditional on Poisson(*θT*) =*m* is again a Gamma distribution with parameters (*k* + *m*, λ + *θ*). This property will be extensively used in the updating of the epoch times occurring during each step of the construction of a particle by the sampler.

### 2.3 Definition of a Particle

Write *S* for the observed SFS. For any fixed *A* = (*A_k_*)_2≥k≥n_, *β* ∈ (–2, ∞) and *θ* > 0, our sampler produces samples from the support of

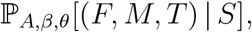

where 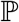_*A,β,θ*_ is the probability measure under which the tree topology follows the incremental Beta-splitting model with parameter *β*, the epoch times are exponentially distributed with parameters *A_k_* and mutations fall on the tree at rate *θ* (an expression for the density of 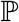_*A,βθ*_ is given in §3.1.2). In particular, our sampler directly produces trees with mutations which are compatible with the observed SFS S, in the sense that no sampled SFS history (*F*, *M*, *T*) satisfies

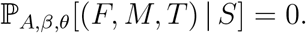

It is devised in a way which maximizes the exploration of the state space of trees with mutations that are compatible with the observed SFS *S*, while biasing the sampling according to the desired balance *β* and epoch times dictated by *A*.

The probability that the sampler produces an SFS history (*F*, *M*, *T*) has density

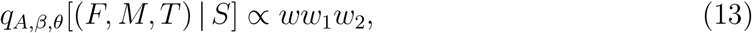

where *w*, *w*_1_ and *w*_2_ are the weights output by the sampler corresponding to *F*, *M* and *T*, respectively. A *particle* refers to such an *S*-compatible SFS history and the corresponding weights, i.e., [(*F*, *M*, *T*), (*w*, *w*_1_ and *w*_2_). Importance sampling estimators are based on a collection of such particles and thus the most basic task of the sampler is the probabilistic construction of a particle. Below we provide an overview of how a particle is constructed incrementally in three stages. The detailed pseudo-code can be found in Section 1 of the Supplementary Material.

### 2.4 The Sampler

The construction of a particle (*F*, *M*, *T*) is an incremental process. It starts from a topology matrix *F* and a mutation matrix *M* whose entries are all zero, a vector of epoch times drawn from the independent exponential *a priori* distributions with rates given by the vector *A*, and the weights *w*, *w*_1_ and *w*_2_ are all set to 1. First, we use *S*_*n*-1_, the number of mutations carried by exactly *n* – 1 individuals in the sample, to force the presence of an edge subtending *n* – 1 leaves if *S*_*n*-1_ > 0. We distribute the *S*_*n*-1_ mutations on the appropriate edges and update the corresponding epoch times. Then, we update the result of this partial construction by distributing the *S*_*n*-2_ mutations carried by *n* – 2 individuals on the edges subtending *n* – 2 leaves (which are forced to exist if *S*_*n*-2_ > 0) and by resampling the corresponding epoch times conditionally on the partial mutation pattern. We then use the mutations carried by *n* – 3 individuals to update the new value of (*F*, *M*, *T*) by possibly inserting some edges subtending *n* – 3 leaves, and so on. Hence, the j-th step of this algorithm considers mutations carried by *n* – *j* individuals and leads to the potential insertion of some (*n* – *j*)-edges (and only such edges) in the partial topology.

More precisely, each of these steps (say, the *j*-th one) goes through three stages (an example with n = 5 is presented below):

i. **Updating the topology:** We update the topology matrix *F* and the weight w obtained at the end of step *j* – 1 based on whether *S*_*n*-*j*_ > 0 (and we should see at least one edge subtending *n* – *j* leaves) or not. We start in epoch 2 and go down the partially constructed topology until there is a possibility for the creation of an (*n* – *j*)-edge. Suppose first that *n* – *j* > n/2, so that there may be at most one such edge in the tree, created by the splitting of a bigger edge. Because this ‘parent’ edge subtends *m* > *n* – *j* leaves, the epoch at which it is split is already encoded in the partially constructed *F* matrix (since the presence or absence of *m*-edges in every epoch is already fully determined for every *m* > *n* – *j*). At the moment when this m-edge splits, we also know that *n* – *j* is the biggest size possible for the largest ‘daughter’ edge (otherwise the creation of an edge subtending more than *n* – *j* leaves at the time of this split would already be recorded). Thus, if *S*_*n*-*j*_ > 0 we force the creation of an (*n* – *j*)-edge at the epoch starting at the split. If *S*_*n*-*j*_ = 0 we have no constraints, and so we decide whether the split gives rise to an (*n* – *j*)-edge or not at random, using the Beta-splitting distribution λ_*m*_,. with parameter *β*, *conditional* on the size of the largest daughter edge being at most *n* – *j*. We also multiply the weight w by the corresponding conditional probability. If the (*n* – *j*)-edge is indeed created, we now have to decide in which epoch it is split and gives rise to two smaller edges. To this end, let us observe that all the other edges present at the same time in the tree are necessarily smaller (recall that *n*–*j* > *n*/2). Consequently, no split has yet been fixed in the remaining epochs until epoch *n*. As in the description of the incremental Beta-splitting model, we carry on going down the tree and decide at the beginning of each epoch *k* to split the (*n* – *j*)-edge with probability (*n* – *j* – 1)/(*n* – *k* + 1), or to keep it with probability (*j* + 2 – *k*)/(*n* – *k* + 1), until the split occurs and the edge disappears from the later epochs (i.e., *F*(*k*, *n* – *j*) = 0 again). Note that the split probability is 1 when *k* = *j* + 2, corresponding to the fact that there can be no (*n* – *j*)-edges in epoch *k* > *j* + 1. The weight *w* is updated accordingly. When *n* – *j* ≤ *n*/2, the procedure is similar but we have to take into account the information present in the partial topology resulting from the previous steps, which could force the creation of an (*n* – *j*)-edge independently of the presence of mutations carried by *n* – *j* individuals in the sample (if the split of an *m*-edge and the subsequent creation of an (*m* – (*n* – *j*))-edge are already encoded in the partial topology, or if *n* is even and a split (*n*/2, *n*/2) is the only remaining option for the first split). The different cases 1 ≤ *j* < *n*/2, *j* = *n*/2 and *n*/2 < *j* ≤ *n* – 3 are handled respectively by the procedures `Sstep, Hstep` and `Lstep`. Eventually, there is no randomness in the insertion of the edges subtending 2 or 1 leaves, which are carried out by the procedures `Twostep` and `Onestep`. All these procedures are listed in Section 1 of the Supplementary Material.
ii. **Distributing the mutations:** The mutations carried by *n* – *j* individuals are placed on the tree. We exploit the information given by the vector *T* produced by the previous steps (i.e., the information obtained from mutations carried by more than *n* – *j* individuals) to give a weight to each epoch containing at least one edge subtending *n* – *j* leaves. Then we distribute the *S*_*n*-*j*_ mutations in a multinomial way, using these weights. The simple idea behind this multinomial scheme is that if mutations fall on the tree like a Poisson point process with fixed rate, then conditionally on there being *S*_*n*-*j*_ mutations carried by *n* – *j* individuals, they are independently and uniformly distributed over the total length of (*n* – *j*)-edges in the tree. Thus, if the state of the epoch time vector just before this step is (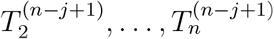), the total length of (*n* – *j*)-edges in any epoch *k* in the current state of the tree is *F*(*k*, *n* – *j*)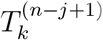, and for each of the *S*_*n*-*j*_ mutations to be placed (independently) on the tree, the probability that it occurs in epoch *k* is

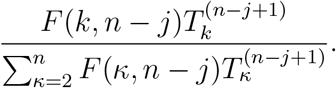

We also update the current value of the weight *w*_1_ by multiplying it by the probability of the mutation placement obtained. Of course, if *S*_*n*-*j*_ = 0 there is nothing to do, even if an (*n* – *j*)-edge exists.
iii. **Updating the epoch times:** We only update the lengths of the epochs containing at least one (*n*–*j*)-edge (since the distribution of mutations gives no new information on the epochs where there are no such edges). To this end, we use the stability property of the Gamma distributions expounded in the previous section: If *T* ∼ *G*(*m*, λ), then *T* conditioned on the event {Poisson(*θT*) = *s*} is a Gamma(*m* + *s*, λ + *θ*) random variable. Using the previous steps and the fact that an exponential distribution with rate *A_k_* is also a *G*(1, *A_k_*) distribution, for every epoch *k* we know that the current value of *T_k_* is a random draw from a *G*(*m*_−_, λ_−_) distribution with

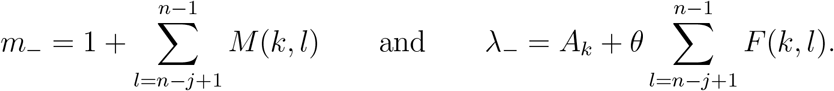 Thus, for every epoch *k* in which an (*n* – *j*)-edge was placed during the first stage, we draw a new value of *T_k_* from a *G*(*m*_+_, λ_+_) distribution with

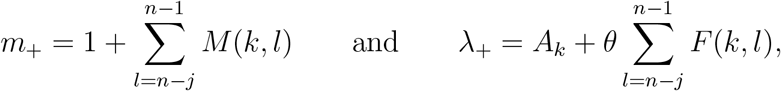

and multiply the weight *w*2 corresponding to this component by the density of the Gamma variable at the value drawn.

MakeHistory listed as Function 1 in Section 1 of the Supplementary Material outputs the desired particle as a list [*F*, *M*, *T*, *w*, *w*_1_ and *w*_2_] when called with the following input arguments: the sample size *n*, the SFS *S*, the scaled mutation rate *θ*, the tree shape parameter *β* and the vector *A* of *a priori* rates for the epoch times. The presence or absence of mutations carried by each number of individuals is encoded in a control sequence *C* = (*C*_1_,…, *C*_*n*–1_), constructed at the beginning of the procedure by setting *C_j_* = 1 if *S_j_* > 0, and *C_j_* = 0 otherwise. The control value *C_j_* = 1 forces the insertion of a first *j*-edge in the tree, after which *C_j_* is set to 0 and the insertion of other *j*-edges remains possible but is not compulsory.

### 2.5 Example

Take *n* = 5 and suppose that the observed SFS is *S* = (4, 0, 1, 0). We give an example of the construction of a particle with *β* = 0 and *A_k_* = *k*(*k* – 1)/2, i.e., the parameters corresponding to the Kingman coalescent. The topology and mutation matrices *F* and *M* are initialized at 0, and the epoch time vector *T* is initialized at a vector of samples from independent random variables with distribution Exp(*k*(*k* – 1)/2). The weights *w*, *w*_1_ and *w*_2_ are initialized at 1. Finally, the control vector is set to be (1, 0,1,0) since there are mutations carried by 1 and 3 individuals, but no mutations are carried by 2 or 4 individuals in the sample. For the sake of clarity, below we do not include the construction of the columns 0 and *n* of *F*, which record respectively the size of the largest daughter edge and the size of the edge split at the beginning of the *k*-th epoch.

#### j = 1 (mutations carried by *n* – 1 = 4 individuals)

*Updating F:* Since *S*_4_ = 0, there are no constraints on the presence of a 4–edge in the tree. Using (9), with probability 1/2 we choose to create a 4–edge in epoch 2, thus setting *F*(2, 4) = 1. We also multiply *w* by 1/2. Epoch 2 being the only epoch in which the presence of a 4-edge is possible, the update of *F* based on *S*_4_ stops here and *F*(*k*, 4) remains equal to 0 for all *k* > 2.

*Updating M:* Since *S*_4_ = 0, there are no mutations to place on the tree and *w*_4_ is not updated.

*Updating T:* Because the (only) 4-edge placed in epoch 2 of the tree carries no mutation, we update *T*_2_ only, by replacing its current value by a sample from a Gamma(1,1 + *θ)* variable (i.e., the law of an Exp(1)-random variable *T*_2_ conditioned on Poisson(*θT*_2_) = 0). We also multiply *w*_2_ by the density of the Gamma distribution at the sampled point.

#### j = 2 (mutations carried by *n* – 2 = 3 individuals)

*Updating F:* In epoch 2, we have already placed a 4-edge and so a β-edge is impossible and *F*(2, 3) remains equal to 0. Next, in epoch 3, since *C*_3_ = 1 the split of the 4-edge in the previous epoch needs to lead to the creation of a β-edge (which otherwise could not appear later in the tree), and so we set *F*(3, 3) = 1 with probability 1 (and we set *C*_3_ = 0 since an edge able to accommodate the β-mutation has been included in the tree, and so the presence of another β-edge is not compulsory). Since a β-edge cannot appear in a later epoch, the update of *F* stops here and *F*(*k*, 3) remains equal to 0 for *k* = 4, 5.

*Updating M:* The only epoch containing a 3-edge is epoch 3, and so the mutation carried by 3 individuals needs to appear in this epoch. We thus set *M*(3, 3) = *S*_3_(= 1) and keep the other *M*(*k*, 3) equal to 0. The weight *w*_1_ is not updated since the chosen mutation placement has probability 1.

*Updating T:* Since epoch 3 is the only epoch containing a 3-edge, *T*_3_ is the only time about which we have some new information for a possible update. As *M*(3, 3) = 1, we update *T*_3_ by sampling a new time from a Gamma(2, 3 + *θ*) distribution and we multiply *w*_2_ by the density of the Gamma distribution at the sampled point.

#### j = 3 (mutations carried by *n* – 3 = 2 individuals)

*Updating F:* In epoch 2 (resp., 3), the presence of the 4-(resp., 3-)edge prevents the presence of a 2-edge, so that *F*(2, 2) = 0 = *F*(3, 2). The 3-edge being the one split between epochs 3 and 4 (since *F*(4, 3) – *F*(3, 3) = –1), it necessarily leads to the creation of a 2-edge in epoch 4. Hence, we set *F*(4, 2) = 1 and we do not need to update *w* since this move has probability 1. The 2-edge is the only possible edge to split in the next (and last) epoch, and so *F*(5, 2) = 0 (and *w* does not need to be updated).

*Updating M*: *S*_2_ = 0, and so there are no mutations carried by two individuals to place in the tree. Consequently, neither *M* nor *w*_1_ is updated.

*Updating T:* The only 2-edge in the tree is that appearing in epoch 4, on which *S*_2_ = 0 mutations are placed, and so the current value of *T*_4_ is replaced by a sample from a Gamma(1,6 + *θ*) random variable and *w*_2_ is multiplied by the value of the density of this Gamma distribution at the sampled point.

#### j = 4 (mutations carried by *n* – 4=1 individuals)

*Updating F:* As for the 2-edges, there is no randomness in the placement of 1-edges in the topology. In epoch 2, the creation of the 4-edge from the split of the ancestral 5-edge imposes that *F*(2,1) = 1. This edge cannot be split further, and so its remains in every later epoch. In epoch 3, the creation of the 3-edge from the split of a 4-edge leads to the creation of a new 1-edge, so that *F*(3,1) = 2. Likewise, *F*(4,1) = 3 and since the 2-edge in epoch 4 is split into two 1-edges, we necessarily have *F*(5,1) = 5. The weight *w* is not updated since all these changes have probability 1.

*Updating M:* For each of the *S*_1_ = 4 mutations carried by one individual, independently of each other, the probability *p_k_* that it is placed in epoch *k* is given by

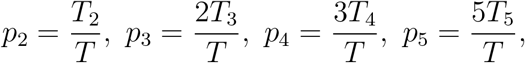

with *T* = *T*_2_ + 2*T*_3_ + 3*T*_4_ + 5*T*_5_. We thus draw the column (*M*(*k*, 1))_2≤*k*≤5_ from a Multinomial(4;*p*_2_,*p*_3_,*p*_4_,*p*_5_) distribution, and we multiply the weight w_2_ by the probability of this draw. For instance, *M*(2,1) = 0, *M*(3,1) = 1, *M*(4,1) = 1 and *M*(5,1) = 2.

*Updating T:* We have placed no mutations carried by one individual on the single 1-edge in epoch 2, and so *T*_2_ is replaced by a sample from a Gamma(1,1 + 2*θ*) r.v. and *w*_2_ is multiplied by the value of the density of the Gamma distribution at the sampled point. Likewise, *T*_3_ is replaced by an independent sample from a Gamma(3, 3 + 3*θ*) variable, *T*_4_ is replaced by a sample from a Gamma(2,6 + 4*θ*) variable and *T*_5_ is resampled from a Gamma(3,10 + 5*θ*) variable. The weight *w*_2_ is also multiplied by the values of densities of these Gamma distributions at the sampled points.

The sampler then outputs the topology and mutation matrices

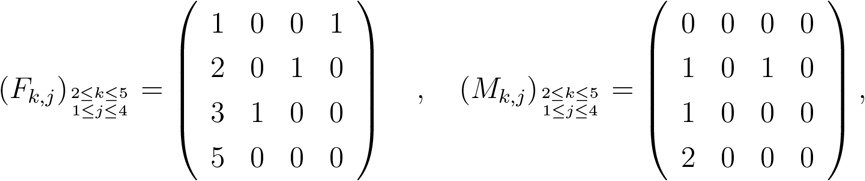

as well as the final state of the epoch time vector *T* and of the weights *w* (= 1/2 in this simple example), *w*_1_ and *w*_2_.

## 3 Simulations and likelihood computations

We explored the properties of our sampler through simulations from the Beta-splitting extensions of two standard models of demography, the exponential growth model and the bottleneck model. Both are recalled and their likelihood functions are given in Section 3 of the Supplementary Material. We also produced data with different *β* parameters in order to test whether our likelihood procedure was able to detect deviations from the Kingman (or equal rates) topology corresponding to *β* = 0. Recall that *β* → – 2 yields more and more imbalanced trees, while *β* → ∞ gives rise to more and more balanced trees. As a sanity check, we also produced simulations with very large mutations rates (yielding of the order of 300 or more SNPs per locus for a sample size of *n* = 15) to check that we were able to recover the true parameters from a reasonable number of loci and particles per locus (60 loci and 200 particles per locus in our simulations, with various samples). These results are not shown here but are available (and can be reproduced) at [SV18].

### 3.1 Likelihood computations

To estimate the likelihood of a given demographic and structural scenario, we use the standard importance sampling methodology [TK10]. It may be improved in different ways, in particular the step-by-step construction of each ‘particle’ may be the basis of an adaptive importance sampling scheme with resampling and partial ‘mutation’ of the particles (see the last sections in [TK10]). We adopt two different approaches, depending on whether the unconditional distribution of genealogical trees is available or not for the class of scenarios considered. In both cases, to produce the genealogies with mutations we need to inform the sampler with a balance parameter *β*, a mutation rate *θ* and a vector of rates for the *a priori* exponential distributions of the epoch times.

The mutation rate and balance parameters, if unknown, may be considered as parameters to be estimated by the procedure. As concerns the vector A giving the rates in the *a priori* distribution of the epoch time vector, if no explicit expressions exist for the class of scenarios considered, the rates can be obtained by a round of simulations of genealogical trees under the scenario considered without conditioning on the observed SFS. In this way, we can estimate the mean epoch times under this scenario and take the inverse of these averages as *A_k_*.

#### 3.1.1 Scenarios for which a likelihood function is available

Suppose that we consider a class of demographic and structural scenarios *ℱ* under which the law **P**_*f*_ of the genealogy of a sample (not conditioned on the observed SFS *S*) is known. In this case, we can evaluate the likelihood of a scenario *f* by sampling m independent particles [(*F_i_*, *M_i_*, *T_i_*), (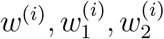)] and by using the fact that for *m* large enough

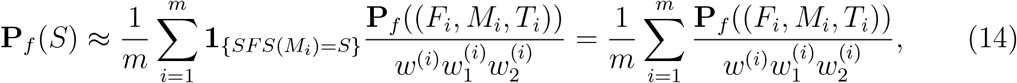

where we abuse notation and write also **P**_*f*_ for the density function of the distribution **P**_*f*_ (observe that the law of the epoch time vector is absolutely continuous with respect to Lebesgue measure). The last equality comes from the fact that the sampler produces particles which are always compatible with the SFS given as an argument. Assuming the infinitely many site model with Poissonian mutation occurring at rate *θ*, we can further decompose the expressions in the numerators as

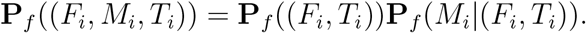

We know the (density of the) law of the genealogical tree (*F*, *T*) under **P**_*f*_, and furthermore the law of *M* knowing (*F*, *T*) is the product over all epochs *k* and numbers of carriers *j* of the probability of that a Poisson(*θ*(*k,j*)*T_k_*) variable is equal to *M*(*k,j*):

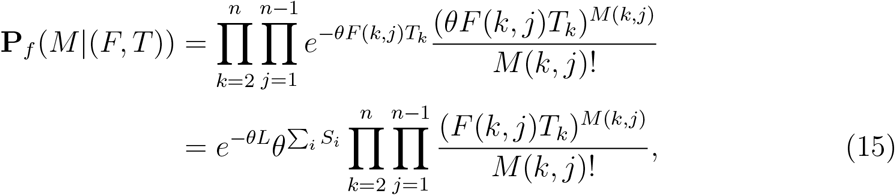

where 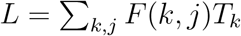 is the total length of the tree. Consequently, every term in (14) can be computed to obtain an approximation to the likelihood of *f* ∈ *F*. Note that *F* does not need to be a parametric family here.

In our simulation studies, we use a version of (14) in which **P**_*f*_ ((*F_i_*, *M_i_*, *T_i_*)) is replaced by **P**_*f*_ ((*F_i_*, *S*, *T_i_*)). This modification was motivated by the fact that for large sample sizes, the product in (15) may have very small values compared to the probability of the SFS knowing the tree, because of the large number of possible mutation placements corresponding to the same SFS. Although we do not have a precise justification for it, the modified likelihood procedure gives very good results in practice.

#### 3.1.2 Scenarios for which a likelihood function is not available

Suppose that we do not have an analytic expression for the (unconditional) distribution of the genealogies under the scenarios of interest. In this case, we can resort to finding the Beta-splitting demographic model with exponential epoch times which best fits the data. Recall our notation 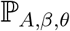 for the probability measure under which (*i*) the genealogy of a sample follows the incremental Beta-splitting model with parameter *β* defined in §2.1, (*ii*) with independent exponentially distributed epoch times whose parameters (or rates) are given by the vector A, and (*iii*) with mutations occurring on the tree at rate *θ*. That is, using (10), the expression for the density of exponential random variables and (15), the density of a given particle (*F*, *M*, *T*) under 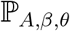 (which we also write 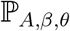 for simplicity) is given by

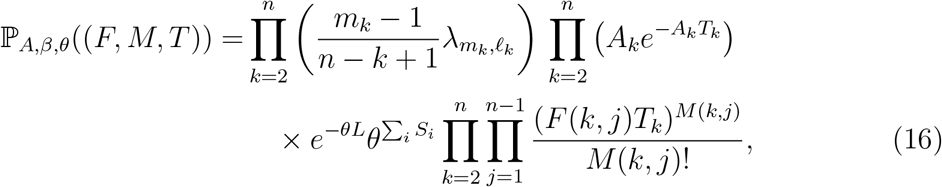

where *L* is the total length of the tree.

Using the same method as in the previous paragraph, we can estimate the likelihood of (*A*, *β*, *θ*) as

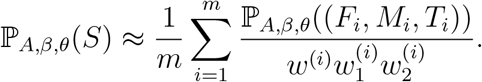

Note that if the law of the genealogy with mutations output by the sampler is close to the distribution 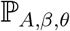 (⋅ | *S*), then we do recover that the importance weights satisfy

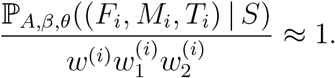

In this case, each ratio 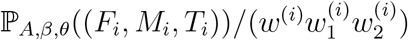 is very close to 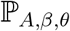(S) and only a small number of particles is sufficient to obtain a precise approximation for 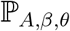(*S*). However, because the construction of a particle (and its weights) is incremental in the number *j* of mutation carriers while the computation of the probability 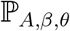((*F*, *M*, *T*)) is incremental in the epoch labels *k*, we do not have an analytic way to check how close the sampler’s distribution is to the conditional law 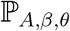 (⋅ | *S*).

In the following sections, we present a few parametric examples of the procedure described above.

### 3.2 Exponential growth model

In Figure 2, we show the likelihood surface for the pair (*θ*, *g*), where *θ* is the scaled per locus mutation rate and *g* is the population growth rate, based on simulated data from the exponential growth model and *β* = 0. The true parameters are *θ* = *ϕ*_1_ = 20, *g* = *ϕ*_2_ = 0 (no growth) and *β* = 0 corresponding to the standard Kingman coalescent. Three independent SFS were simulated, corresponding to 3 independent loci and sample size *n* = 30. The 2-dimensional grid over *θ* and *g* is explored with *β* fixed at 0 via a quasi Monte Carlo scheme, and for each SFS and each pair (*θ*,*g*), the evaluation of the likelihood is based on 1000 particles. The top two and bottom-left subplots show the likelihood surfaces obtained using only a single locus (labelled 0, 1 and 2), the bottom right subplot shows the product of the likelihoods for all three loci.

**Figure 2:**
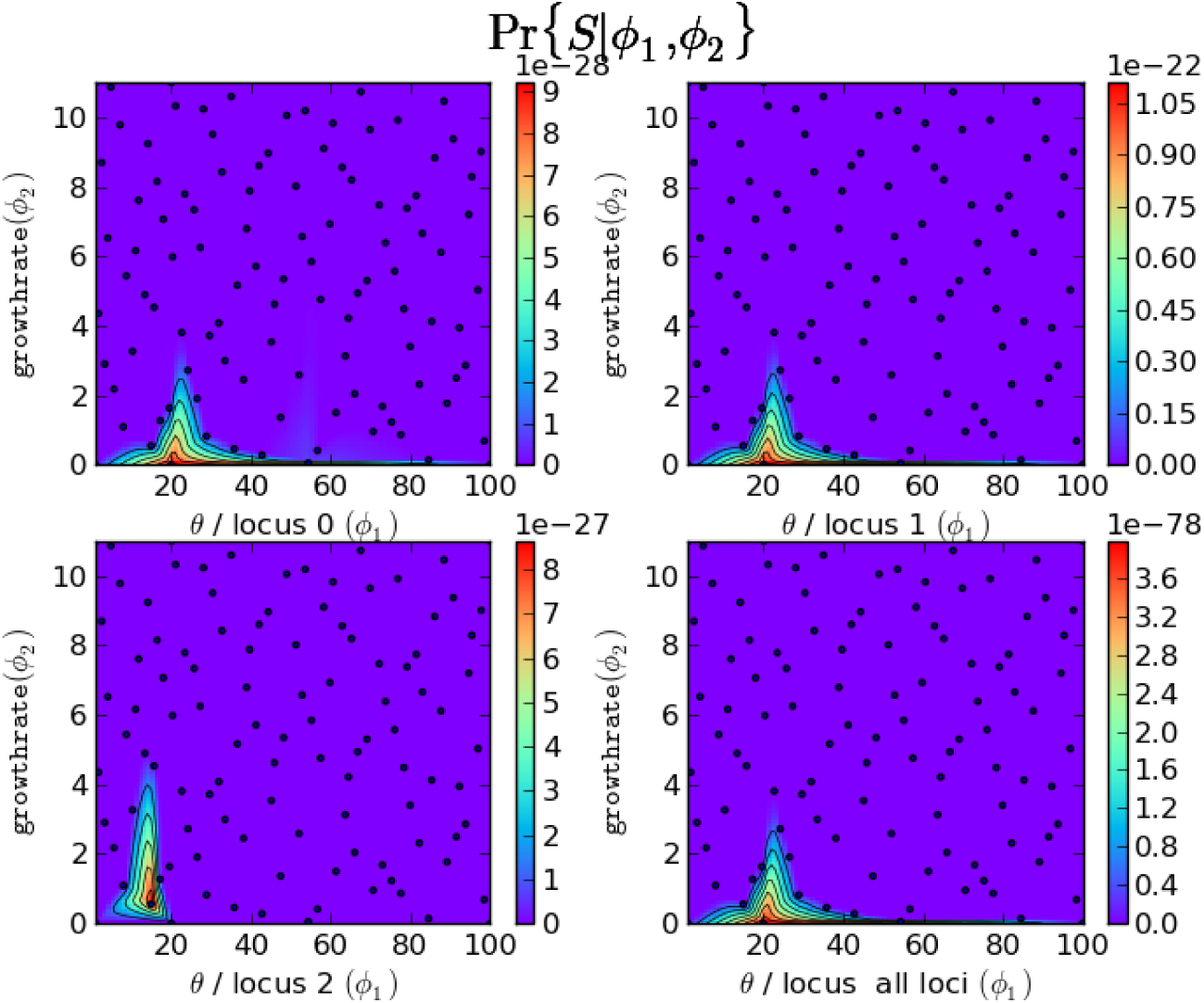
Likelihood surfaces of *θ* = *ϕ*_1_, the per locus scaled mutation rate, and population growth rate *g* = *ϕ*_2_ for three loci. True parameters are *θ* = 20, *g* = 0 and *β* = 0 with a sample size of 30. The black dots shows the points in the parameter space sampled by the quasi-Monte Carlo procedure, except in the region of higher likelihood where the density of sampled parameters is high. The likelihood calculations are based on 1000 particles per parameter per locus.

Of course each SFS corresponds to a single realization of the genealogy with Poissonian mutation of the sample, and so we expect the maximum likelihood estimator to improve with the number of independent loci considered. This is indeed the case in another simulation study involving 100 loci over a coarse proof-of-concept grid, as shown in Table 1, in which we also make the parameter β vary. Here the likelihood estimates are based only on 100 particles, which in general should be considered as the minimal number of particles that should be sampled per parameter per locus to ensure a reasonable precision of the estimation via the law of large numbers. However, increasing the number of loci or the number of particles (*F*, *M*, *T*) sampled to compute the likelihood corresponding to a given locus has a computational cost, and it is therefore important to assess the capabilities of our procedure with reasonable numbers of loci and particles per locus. Table 1 suggests that the number of loci need not be very large for the parameter estimates to be reliable.

**Table 1:** Log-likelihood (log-lkh) of parameters *g* (growth rate), *θ* (scaled per-locus mutation rate) and *β* (balance parameter) under the Beta-splitting model with growth. True parameters are *g* = 0, *θ* =10 and *β* = 0, the sample size is *n* =15 and 100 independent SFS have been generated. The mean number of SNPs per locus is 70.05. For each SFS and triplets of parameters, the likelihood estimate is based on 100 particles. The partial log-likelihood is obtained by considering only the first 30 loci, the full log-likelihood uses all 100 SFS. In both cases, the maximum likelihood estimate is obtained for 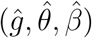 = (0,10,0). The parameter points in the Cartesian grid for (*g*, *θ*, *β*) ∈ {0,1,10} x {1,10,100} x {-1,0,10} with likelihood estimates of –∞ in double precision are not shown.

| $g$ | $\theta$ | $\beta$ | Partial log-lkh | Full log-lkh |
| --- | --- | --- | --- | --- |
| 0 | 1 | -1 | -2337.0 | -7667.8 |
| 0 | 10 | -1 | -1576.6 | -5226.2 |
| 0 | 100 | -1 | -3065.1 | -10086.1 |
| 0 | 1 | 0 | -2345.4 | -7599.9 |
| 0 | 10 | 0 | **-1569.6** | **-5110.1** |
| 0 | 100 | 0 | -3031.3 | -10196.8 |
| 0 | 1 | 10 | -2340.0 | -7643.8 |
| 0 | 10 | 10 | -1581.8 | -5324.1 |
| 0 | 100 | 10 | -3146.0 | -10321.1 |
| 1 | 10 | -1 | -1817.2 | -5909.2 |
| 1 | 100 | -1 | -3047.5 | -10086.5 |
| 1 | 10 | 0 | -1732.8 | -5786.1 |
| 1 | 100 | 0 | -3036.4 | -10154.4 |
| 1 | 10 | 10 | -1812.2 | -5853.3 |
| 1 | 100 | 10 | -3157.5 | -10255.5 |
| 10 | 100 | -1 | -3031.2 | -9975.8 |
| 10 | 100 | 0 | -3019.1 | -10014.9 |
| 10 | 100 | 10 | -3108.1 | -10230.8 |

Table 2 shows another simulation study when the growth rate is non-zero. Furthermore, small sample sizes are sufficient to detect population growth in the exponential growth model. Indeed, for moderate to large sample sizes the first coalescence events happen very quickly, and at that time the population size has not sufficiently decreased (backwards in time) to have a strong impact on the total edge length in these epochs, and thus on the distribution of mutations on which our procedure is based. In Table 2, we only consider samples of size 2. The parameter *β* does not play a role in this case and so we set it to 0 in all likelihood calculations. In this case, 30 loci and 1000 particles per locus per SFS are sufficient to detect a deviation from the hypothesis *g* = 0, and to correctly estimate the true value of *g* and *θ* in the short list provided as an example.

**Table 2:** Log-likelihood (log-lkh) of parameters *g* and *θ* under the Beta-splitting model with exponential population growth. Here the true parameter values are *g* = 10, *θ* = 10 and *β* = 0. For a sample of size 2, 100 independent SFS were produced (the mean number of SNPs per locus is 3.82), but only 30 of them are used to compute the approximate log-likelihoods. 1000 particles are produced per SFS and per pair (*g*,*θ*). The most likely parameter values in this short list are 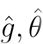 = (10,10). The parameter *β* is not estimated here as a sample of size 2 brings no information on the balance of the genealogy.

| $g$ | $\theta$ | Partial log-lkh |
| --- | --- | --- |
| 0 | 1 | -86.9 |
| 0 | 10 | -97.8 |
| 0 | 100 | -159.8 |
| 1 | 1 | -115.5 |
| 1 | 10 | -91.7 |
| 1 | 100 | -159.0 |
| 10 | 1 | -429.9 |
| 10 | 10 | **-71.7** |
| 10 | 100 | -151.7 |

In order to assess the reliability of our maximum likelihood inference procedure, we simulated a set of 100 independent SFS under the standard Kingman coalescent scenario (*β* = 0 = *g*, *θ* = 10) with sample size *n* = 4. We then produced 10 estimates for the likelihood of the true parameter, and for that of a close but erroneous parameter (*β*, *θ*, *g*) = (0,10,1), using different numbers of particles per locus in the likelihood estimation. As shown in Table 3, as the number of particles per locus increases, the distribution of the log-likelihood estimate for each parameter becomes more and more concentrated. Furthermore, the log-likelihood estimates of the true parameter (0, 10, 0) become significantly larger than those of the erroneous parameter (0,10,1). Due to the order of magnitude difference in log-likelihood values between distant parameter values, we found that even 20 particles are often sufficient to quickly explore the relative log-likelihoods over a coarse grid of parameters. A more careful exploration of the parameter space will require a larger number of particles per locus (about 100 particles at least) in general.

**Table 3:** Concentration of the log-likelihood estimates as the number of particles per locus used in the procedure increases. We simulated 100 independent SFS under the standard Kingman coalescent scenario (true parameter (*β*, *θ*, *g*) = (0,10, 0)). Based on this series of SFS, we produced 10 estimates of the log-likelihood of the true parameter and of some close but erroneous parameter (0,10,1), using different numbers of particles per locus (per parameter) in the importance sampling procedure. For each parameter value and each number of particles per locus, we report the mean and standard error of the 10 replicates.

|  | Nb particles | 1 | 10 | 100 | 500 | 1000 |
| --- | --- | --- | --- | --- | --- | --- |
| (0, 10, 0) | mean log-lkh | -1126.7 | -649.2 | -516.8 | -454.5 | -425.1 |
|  | std. err. | 13.3 | 10.5 | 4.0 | 3.3 | 2.6 |
| (0, 10, 1) | mean log-lkh | -1599.5 | -903.9 | -690.7 | -613.8 | -583.7 |
|  | std. err. | 35.3 | 10.6 | 5.9 | 4.6 | 3.4 |

Regarding the estimation of the tree shape parameter *β*, sufficiently large deviations from the Kingman case *β* = 0, such as *β* = −1.9 or *β* = 50 are easily detected by our procedure. Milder deviations of *β* from 0 such as –1 or 10 can be detected with 30 loci with sample size of 15 as shown in Table 4. Distinguishing smaller deviations of *β* from 0 is slightly more delicate as it seems that the law of the topology varies slowly with the parameter *β*. Of course the larger the sample size, the more splits there are in the tree to estimate the different transition probabilities. However, our simulation studies suggest that a sample size of *n* =10 is already sufficient to obtain close estimates, as long as the number of particles per locus per parameter value is sufficiently large (from 100 to 1000, for example).

**Table 4:**
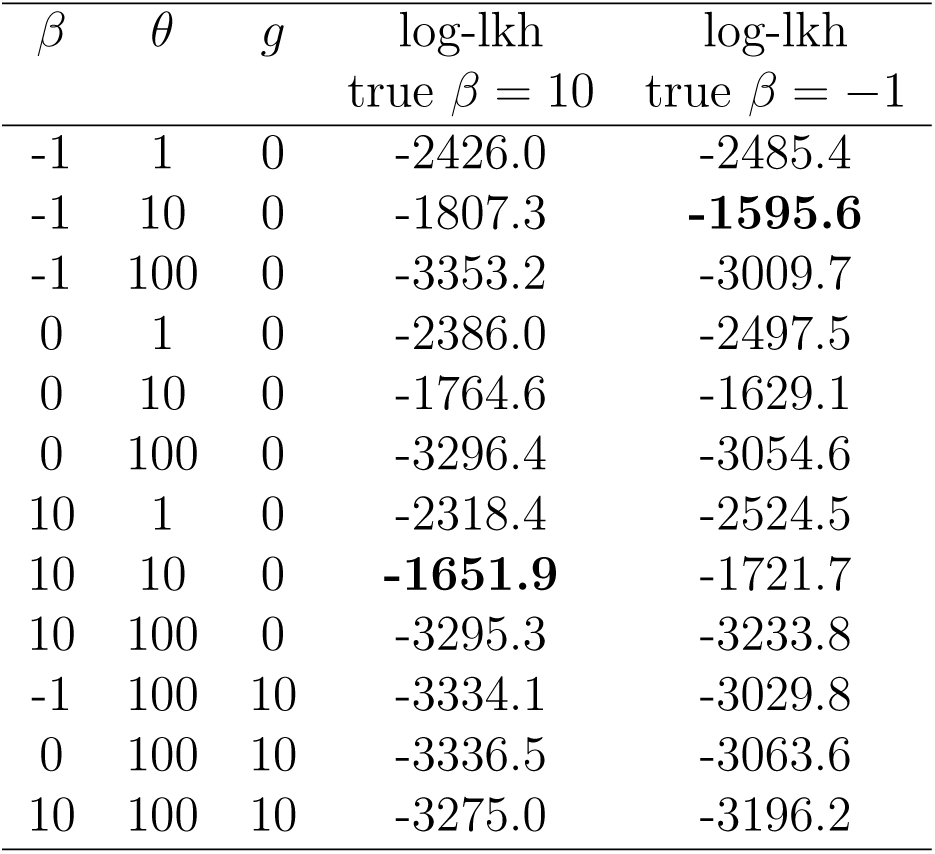
Log-likelihood (log-lkh) of parameters *β*, *θ* and *g* under the Beta-splitting model with exponential population growth. Here the true parameter values are *g* = 0, *θ* =10 with (i) an unbalanced *β* = –1 and (ii) a more balanced *β* = 10. SFS at 30 independent loci were simulated for a sample size of 15. The log-lkh was only based on 20 particles per SFS per triplet (*β*, *θ*, *g*) ∈ { –1,0,10} x {1,10,100} x {0,10}. The procedure detects the true parameters corresponding to the maximum log-lkh values in bold. Rows with log likelihood estimates of –∞ in double precision are not shown.

### 3.3 Integration with ms and systematic parameter search

We integrated Hudson’s `ms` program [Hud02] with a minor modification to output SFS in order to validate against our simulators and estimators. For example, we were able to estimate the true parameters (same parameter setting as in Table 1) when the SFS were directly simulated from `ms` as opposed to our simulators. This integration with `ms` will allow one to simulate SFS data under rather complex demographic and structural scenarios and simply fit this data to a Beta-splitting demographic model as done in §3.5.

We also confirmed that the true parameters are recovered through a stochastic global optimization algorithm that evolves a population of parameters towards the MLE that concentrates about the true parameters [SP97], albeit at a larger computational cost. These results are not given here as they are fully reproducible from [SV18]. Having demonstrated that systematic parameter searches can be done (especially with optimized version of our code and/or with more computational resources), we henceforth resort to coarse grids of parameter values to quickly illustrate our likelihood computations with our proof-of-concept code under different simulation scenarios.

### 3.4 The bottleneck model

We also investigated parameter estimation in the bottleneck model presented in Section 3 of the Supplementary Material. The results are not shown but can be found at [SV18]. We simulated data corresponding to a recent mild bottleneck, starting at *a* = 0.05, ending at *b* = 0.15, with *N*_0_ = 1 and a reduction in population size of *ε* = 0.01. To reduce the number of parameters, we assumed that the scaled mutation rate *θ* and the length of the bottleneck *b* – *a* was known. The only parameters to reconstruct were then *a* and *ε*.

As in the exponential growth model, small sample sizes (*n* = 3) are sufficient to reconstruct the two parameters as long as the number of loci and the number of particles per locus per set of parameters is sufficiently large (30 or 100 loci with 500 or 1000 particles per locus, for an average number of SNPs per locus of the order of 10). Nonetheless, while the starting time of the bottleneck is always well-reconstructed, the population reduction *ε* tends to be overestimated in general (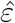 = 0.1), even assuming a larger number of SNPs per locus. Note however that as we increase the average number of SNPs per locus, the likelihood surface becomes more and more peaked. Increasing the sample size to 10 or 20 does not seem to improve the precision of the procedure, probably because the high variability in the topology of larger trees is not compensated by the few additional mutations appearing during the bottleneck and carried by larger numbers of individuals in the sample.

### 3.5 Fitting to the simplest Beta-splitting demographic model

Consider a rather complex historical scenario of a stepping-stone model with a recent barrier (as given in Fig. 3 of the documentation for `ms` [Hud02] available from https://uchicago.box.com/s/l3e5uf13tikfjm7e1il1eujitlsjdx13). There are six subpopulations that exchange migrants in a stepping-stone model. At a time *T* = 2 time units in the past a barrier to gene flow arose, such that no further gene flow occurs between subpopulation 3 and subpopulation 4. We quote the exact command we used in our simulation with explanation quoted directly from the `ms` documentation for concreteness.

`ms 15 100 ‒t 10.0 ‒I 6 0 7 0 0 8 0 ‒m 1 2 2.5 ‒m 2 1 2.5 ‒m 2 3 2.5 ‒m3 2 2.5 ‒m4 5 2.5 ‒m5 4 2.5 ‒m5 6 2.5 ‒m6 5 2.5 ‒em 2.034 2.5 ‒em 2.0 4 3 2.5`

The phrase, -I 6 0 7 0 0 8 0, sets up 6 subpopulations with zero migration rate between them and establishes that a sample of size 7 is taken from subpopulation 2 and a sample of size 8 is taken from subpopulation 5. (In the output the first 7 haplotypes are from subpopulation 2 and the next 8 are from subpopulation 5. The -m commands set up migration, 4Nm = 2.5, between the neighboring subpopulations (except between subpopulation 3 and 4). The -em commands modify the migration matrix at time 2.0 in the past such that pastward of this time, migration at rate 4Nm = 2.5 occurs also between subpopulation 3 and 4.

No analytical likelihood expression is available for this complex historical scenario. We can now fit the 100 SFS loci to the simplest Beta-splitting model with constant population size. The log-likelihood is estimated over a coarse grid of parameters over *β* and *θ* using 1000 particles per locus. The maximum log-likelihood value corresponded to more balanced topologies with 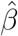 = 5 due to the sampling scheme and the population structure and larger value of *θ* due to longer time to coalescence caused by the barrier with *θ* = 100. Note that when *β* is set to 0 the most likely *θ* is proportional to the product of the mutation rate per locus and the effective population size *N_e_* under the standard Kingman coalescent. Our simplest Beta-splitting model considered here adds an additional tree-balance parameter to the classical setting with constant population size (for details see [SV18]) and is over 60 log-likelihood units better than the model with *β* = 0 and *θ* = 100. We purposely keep the demography of the fitted β-splitting model as simple as possible here to illustrate that some population histories involving complex population structures can be explained significantly by a single tree-balance parameter.

## 4 Discussion

The procedure developed in this work exploits the huge reduction in the size of the hidden space of genealogical trees to explore when focusing on the optimal tree resolution that fully characterizes the law of the SFS. Furthermore, our sampler produces only tree topologies which are compatible with the observed SFS. This double optimization enables us to compute approximate likelihoods for the parameters describing the demographic history of the population, as well as a parameter *β* measuring the typical balance of the genealogy of a sample, at a drastically reduced cost compared to procedures based on the leaf-labelled Kingman coalescent (even with our non-optimized ‘proof of concept’ code). Because the population demographic and structural parameters are shared across (neutral) loci, the per-locus approximate likelihood functions of several hundreds of independent loci can then be combined to bypass the idiosyncratic genealogical history of a single (or a few) loci. For the same reason, dissonant parameter estimates at some loci may enable us to detect outliers, subject to natural selection for example. At the moment, the inference of the parameter *β* can mainly serve to detect deviations from the assumptions of panmixia and neutrality at the basis of the Kingman coalescent model. A thorough investigation of the effect of different kinds of population structure on the topology of the genealogical tree of a sample could for instance lead to a new and simplified criterion for model selection.

The next step is to develop this procedure into an optimized code, enabling us to perform a thorough quantitative comparison with the existing methods for reconstructing demographic parameters. However, we stress again that one of the novelties of this approach is that it is able to compute (approximate) full likelihoods for SFS at nonrecombining loci, which none of the existing methods are able to do (except [BFL15] for very small sample sizes).

### 4.1 Generalizations

Our inference procedure is flexible and could be generalized in many ways. For instance, the mutation rates could differ between loci to accommodate a potential inhomogeneity in the locus lengths or in the mutation rates along the genome. In addition, because the sampler constructs the compatible SFS histories in an incremental way, we may stop the construction at a step that uses mutations carried by *j* > 1 individuals, for instance if we are only interested in the not-too-recent history of the population. This incremental construction may also lie at the basis of an adaptive exploration of the space of compatible topologies via more sophisticated sequential particle filtering schemes involving genealogical and interacting systems [DM04]. To make the semi-parametric family of Beta-splitting models richer and fit more complex tree topological distributions, we can allow *β*’s to change at each epoch or be drawn from a distribution with minimal change to the sampler, provided we have a much larger number of SFS loci.

### 4.2 Filtering out non-recombining loci

Our method requires an important pre-processing step to find loci as contiguous blocks of segregating sites that are free of intra-locus recombination. One could use for example the simple four-gamete test [HK85], or more complex methods for detecting blocks of loci that are free of intra-locus recombination (for eg. [Pos02]). Such filtered loci can then be summarized into SFS and fed into our pipeline for inference. It is important to note that poorly filtered loci with high levels of recombination will tend to give the topological signal of highly unbalanced tree topologies with a local MLE of *β* close to −2. This is because the fully unbalanced tree is compatible with an SFS that has a positive count at every frequency, i.e. contains singleton, doubleton, …, *i*-ton, …, and (*n* – 1)-ton mutations.

### 4.3 Scalable computing framework

Apache Spark, a unified engine for big data processing [ZXW+16], and the ADAM module [MNH+13] for population genomics in particular, are ideal frameworks for deploying the algorithms developed here for real-world applications at the genomic scale. Rewriting our sageMath/Python codes [SV18] in Scala will allow for Spark transformations and actions of our algorithms in conjunction with the ETL methods already available in ADAM. Such an undertaking is beyond the scope and resources of this study and we hope that others may pursue such possibilities.

### 4.4 Adding BIM resolution via gene trees

In this work we restricted ourselves to the information in the SFS. Adding additional information can be done systematically using the *partially ordered graph of coalescent experiments* of [STH+11], say from the full binary incidence matrix of mutational patterns across sites and individual sequences, via the haplotype frequencies, and can significantly improve our estimators (see [PVWR18]).

## Data accessibility

The code developed for this work is given in the Supplementary Material. It is also publicly shared at https://cocalc.com/projects/ac7f397f-eab9-45fc-9278-f486af09ca55/files/FullLikelihoodInferenceSFS.sagews

